# The spatial and metabolic basis of colony size variation

**DOI:** 10.1101/111153

**Authors:** Jeremy Chacón, Wolfram Möbius, William Harcombe

## Abstract

Spatial structure impacts microbial growth and interactions, with ecological and evolutionary consequences. It is therefore important to quantitatively understand how spatial proximity affects interactions in different environments. We test how proximity influences colony size when either *Escherichia coli* or *Salmonella enterica* are grown on different carbon sources. The importance of colony location changes with species and carbon source. Spatially-explicit, genome-scale metabolic modeling predicts colony size variation, supporting the hypothesis that metabolic mechanisms and diffusion are sufficient to explain the majority of observed variation. Geometrically, individual colony sizes are best predicted by Voronoi diagrams, which identify the territory that is closest to each colony. This means that relative colony growth is largely independent of the distance to colonies beyond those that set territory boundaries. Further, the effect of location increases when colonies take-up resource quickly relative to the diffusion of limiting resources. These analyses made it apparent that the importance of location was smaller than expected for experiments with colonies growing on sugars. The accumulation of toxic byproducts appears to limit the growth of large colonies and reduce variation in colony size. Our work provides an experimentally and theoretically grounded understanding of how location interacts with metabolism and diffusion to influence microbial interactions.

## Introduction

Microbial interactions help determine ecosystem functions, from global nutrient cycling to human health(Cho & Blaser 2012; Arrigo 2005). Spatial structure mediates microbial interactions (Connell et al. 2014), however, the relationship between proximity and strength of interaction remains unclear (Nadell et al. 2016). Quantifying, and being able to predict, the effect of location on microbial interactions is critical for understanding processes from the community evolutionary-ecology of microbial ecosystems to emergent functions such as human health.

Spatial structure modulates the resource competition that shapes microbial communities (Mitri & Foster 2013, Foster & Bell 2012, Stacey et al 2016). The relative strength of competition influences community assembly (David et al. 2015) and stability (Shade et al. 2012), in addition to shaping the selection on microbial traits (Gerardin et al. 2016). Spatial structure alters the scope of competition (Mitri & Foster 2013). In agitated liquid environments all cells tend to have equal access to resources and interactions are global. In contrast, in structured environments, cells interact more strongly with neighbors than with distant individuals. This localizing effect of spatial structure has been repeatedly shown to influence the outcomes of microbial evo-ecological experiments (Gerardin et al. 2016; Mitri et al. 2015; Harcombe et al. 2014; Allen et al. 2013; Allison 2005; Kim et al. 2008; Hansen et al. 2007; Kerr et al. 2002; Greig & Travisano 2008; Dechesne et al. 2008; Chao & Levin 1981; Penn et al. 2012; Gralka et al. 2016).

The specific location of bacteria in spatially-structured environments matters. Within a biofilm or colony, bacteria at the edge have smaller local density and grow faster than those in the center (Persat et al. 2015; Gandhi et al. 2016; Nadell et al. 2016; Pirt 1967), which can segregate competing genotypes (Korolev et al. 2012; Mitri et al. 2015; Hallatschek &Nelson 2010; Hallatschek et al. 2007; Momeni et al. 2014). Between-colony interactions are also influenced by colonies’ locations. A competing colony’s effect is magnified if it is located between a focal colony and a nutrient source(Harcombe et al. 2014). Further, the coexistence of competing genotypes can be highly sensitive to the distance between colonies (Gerardin et al. 2016; Kim et al. 2008). Finally, hints of the importance of location are also being detected in complex natural ecosystems, such as in the microbiome. For example, changes in the spacing between *Aggregatibacter actinomycetemcomitans* and *Streptococcus gordonii* determine virulence in oral abscesses(Stacy et al. 2014).

While it is known that location matters, we lack a rigorous framework for understanding and predicting the impact of location on interactions. Interaction strength can be a function of distance(Kim et al. 2008), but by what distance-based measurement? The distance to the closest competitor, a function of all competitor distances, or a measurement of how competitors divide the available territory? Ecologists often use distance metrics to explain variance in plant growth(Tome & Burkhart 1989), and a linearly-weighted distance model captured a decline in bacterial colony size due to crowding (Guillier et al. 2006). In contrast, Voronoi diagrams, which measure the territory that is closer to a focal colony than any other colonies(Okabe et al. 2000), have been used to investigate pattern formation as bacteria cover a surface(Lloyd & Allen 2015). To date there has not been a rigorous test of the ability of different geometric models to explain variance in colony size.

In addition to this geometric description, the question arises what minimal biophysical model can predict the location-based effects on colony growth in variable environments. Microbes typically interact through the chemical compounds that they consume and excrete(Germerodt et al. 2016; Hibbing et al. 2010). Does accounting for metabolism and diffusion suffice to predict the variation in colony growth? Genome-scale metabolic models and flux balance analysis can quantitatively predict the metabolites that microbes consume and excrete, and therefore can predict the ecological interactions that emerge from intracellular mechanisms (Orth et al. 2010; Mahadevan et al. 2002; Harcombe et al. 2014). Further, diffusion of biomass and nutrients can be incorporated to predict system dynamics in structured environments (Harcombe et al. 2014). We therefore can test whether colony variation is purely a function of metabolism and diffusion by comparing computational predictions against experimental observations. If factors such as toxicity, signals or stochastic differences in lag time drive colony variation then the model, which does not take these effects into account, will do a poor job. Determining the extent to which metabolic mechanisms drive spatial effects will be critical for predicting growth in complex natural settings.

Here we investigated how location influences interactions in arguably the simplest scenario - monocultures grown on homogeneous surfaces. We plated monocultures of either *Escherichia coli* or *Salmonella enterica* on various media and used high-resolution scanners to investigate the size of colonies and the associated variance within each plate. We then used simulations and geometric descriptions of varying complexity to determine how much of the colony variation could be explained by metabolic mechanisms, what aspect of location best explained variation in growth, and how variation was influenced by changes in either nutrient uptake or diffusion. Finally, we investigated one case in which variation differed from expectation and suggest that this deviation was caused by byproduct toxicity. Our work provides a quantitative framework for understanding and predicting the effect of location on colony growth which is independent of both microscopic parameters and knowledge of the positions of the non-nearest neighbors.

## Methods

### Strains and media

We used cells of either *Salmonella enterica* subsp. *Typhimurium* LT2 or *Escherichia coli* K12-MG1655 as our model species. In the genome-scale metabolic modeling, these strains were represented by the iRR_1083 model (Raghunathan et al. 2009) and the ijo_1366 model (Orth et al. 2011), respectively. The Petri dish experiments either used LB media (10g/L tryptone, 10g/L NaCl, 5g/L yeast extract) or a modified Hypho minimal media (7.26mM K2HPO4, 0.88 mM NaH2PO4, 3.78 mM [NH_4_]_2_SO_4_, 0.41 mM MgSO4, 1 ml of a metal mix (Delaney et al. 2013)), with either glucose (8.34 mM), citrate (5.10 mM), lactose (4.17 mM), or acetate (12.5 mM) as the limiting resource. All Petri dishes had 1% agar. Dishes were left open under UV for 30 minutes.

For Petri dish experiments, cells from a 1-day old colony were used. After spreading cells onto a Petri dish, a thin piece of black plastic was placed within the upper lid of the dish to improve contrast and reduce reflections from the Petri dish lid. Petri dishes were then placed onto a Canon Perfection V600 scanner agar side down and a 600dpi image was scanned every 20 minutes for almost 150 hours. Each treatment (a unique combination of a species and a media type) was repeated 4-8 times.

### Image Analysis

Tracking of colony areas over time was performed in three main steps using custom software in Matlab (Supplementary material). First, the initial colony locations were detected. Then, the final colony areas were measured and merged colonies were separated (Fig. 1A). Finally, using the outlines of the final colony areas, the area of each colony was tracked through time. We measured the growth rate of colonies on Petri dishes via the diameter of a hypothetical circular colony with same area(Wimpenny 1979). The growth rate was calculated by regressing diameter over time for the first four hours (12 frames) once growth was detected.

**Figure 1:**
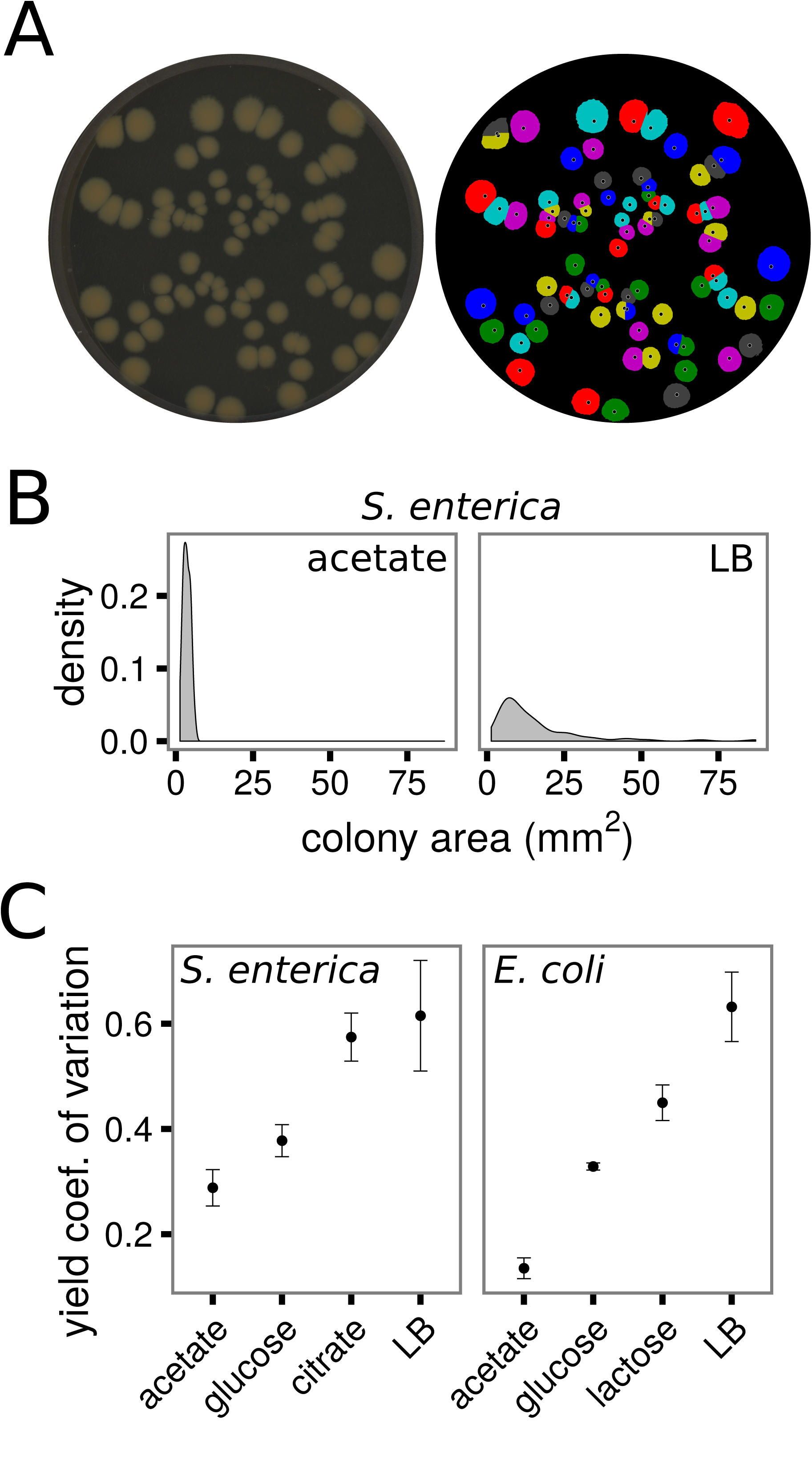
The variance in colony yields depends on the species and environment. A) A snapshot of *S. enterica* colonies on LB media (left) and the yields (areas) of those colonies determined by automated image analysis (right). B) Density distributions of *S. enterica* colony yields grown on acetate or LB. C) The coefficient of variation of colony yields for each species on each carbon course. Error bars are standard error of mean.

### Computational Modeling

All simulations were run in COMETS, a platform developed to do spatially-explicit dynamic flux-balance analysis simulations with an arbitrary number of species (Harcombe et al. 2014). Simulations were carried out using the University of Minnesota Supercomputing Institute’s Mesabi cluster. COMETS uses dynamic flux balance analysis to predict bacterial growth over time by maximizing the amount of biomass which can be produced by a species’ metabolic model given the resources in the local environment (Mahadevan et al. 2002). Michaelis-Menten kinetics constrain resource uptake, with the maximum set by *v_max_* and the concentration at which uptake is at half-maximum set by *k_m_.* In addition, we ran simplified reaction-diffusion simulations in COMETS by using a metabolic model with a single carbon uptake reaction, paired with a reaction that converted all intracellular carbon directly to biomass. In this simplified case uptake kinetics exactly match growth kinetics.

COMETS simulates growth in spatially-explicit environments by solving dynamic flux balance analyses in discrete “boxes” within a lattice at each time step, and then allowing diffusion of the resources and biomass among boxes between time steps. Each box can contain different amounts of biomass or resources, which determine the outcome of the dynamic flux balance analysis. Biomass and resources each diffuse to neighboring boxes with specific diffusion coefficients. The time-step and box sizes we used were small enough to not cause a change in results if we reduced these sizes further.

The starting conditions, and length of time the simulations ran, depended on the question of interest. For the simulations in figures 3 and 4 the “world” was a 5 cm × 5 cm square, into which 60 colonies were seeded at random locations, with initial biomasses of 1e-10 grams. Resources were distributed uniformly at a concentration of 1e-6 mmol per box for the limiting resource, and at essentially infinite abundance for non-limiting resources. These simulations were run until resources were fully consumed. The genome-scale metabolic model simulations, Fig. 2, were conducted in circular environments that were 90 mm in diameter to mimic the experimental conditions. Biomass was seeded at the same location as observed in experiments, and the concentration of the limiting resource also matched the laboratory conditions. Genome-scale simulations were run for equal lengths of time as the laboratory experiments. Other simulation parameters are provided in Table 1.

**Figure 2:**
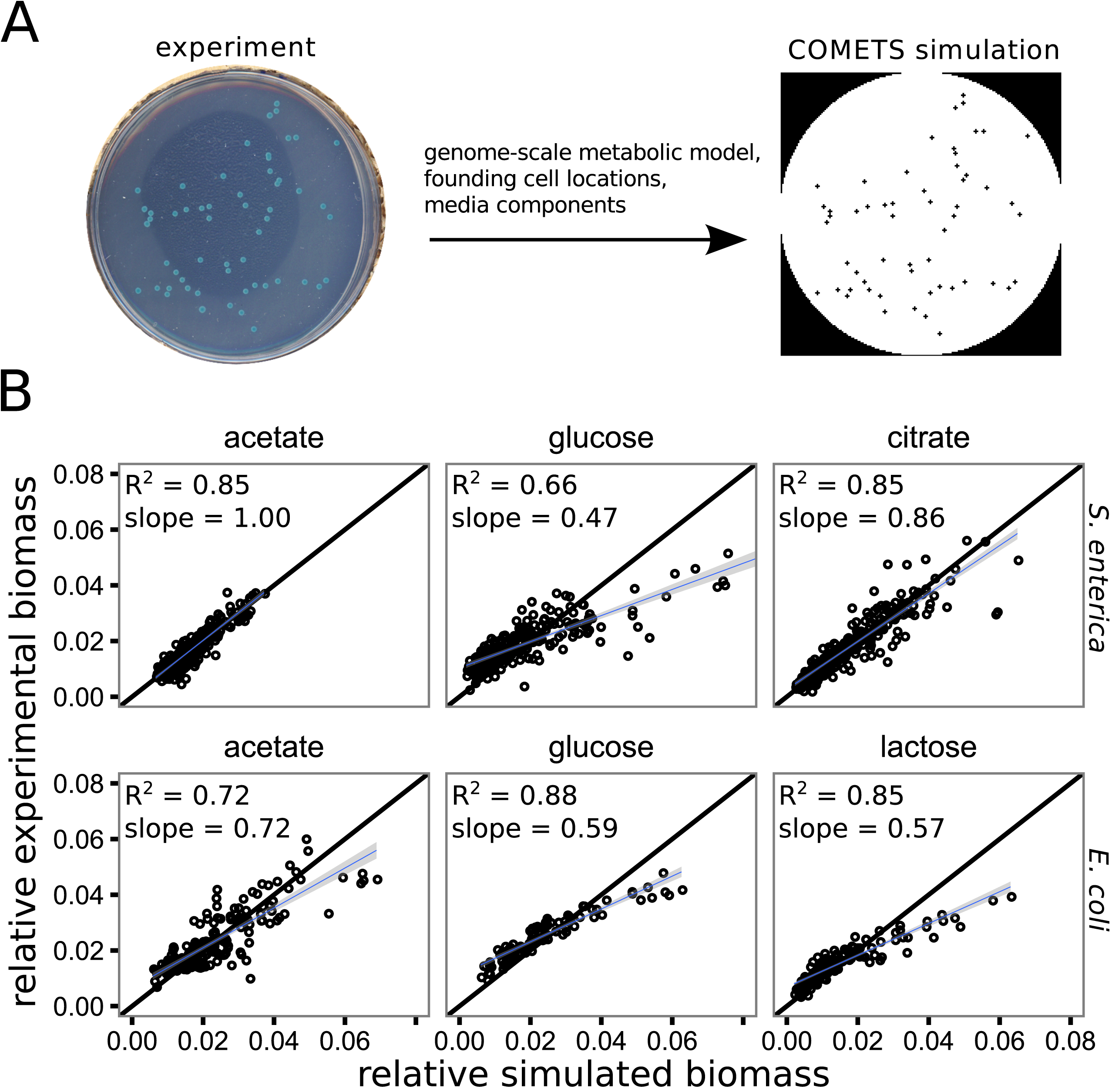
Genome-scale metabolic modeling recapitulates the variance in colony yields. A) We used genome-scale metabolic modeling in the COMETS platform to test the mechanisms generating the observed variance in colony yields. The relevant genome-scale metabolic model was seeded into an environment at the sites from which colonies initiated in experiments. Dynamic flux-balance-analysis calculations and subsequent metabolite and biomass diffusion were carried out in discrete time steps for a duration mimicking the laboratory experiments. B) A comparison of the relative colony yields measured in experiments (y axis) to the relative colony yields predicted by the COMETS simulations (x axis) for the treatments accessible with COMETs (defined media, i.e. not LB). Each facet contains data from 4-8 Petri dish experiments / simulations. The black line has slope = 1 and intercept = 0, while the blue lines and surrounding grey are linear regression lines with standard error. A high R^2^ suggests that the relative spatial effects are captured by the model, while a slope close to 1 suggests an accurate prediction of amount of variance in colony yield.

**Table 1:** Baseline COMETS parameters used for simulations

| Parameter | value | Unit, description |
| --- | --- | --- |
| spaceWidth | 0.05 | Side length of a box, in cm |
| deathRate | 0 | Constant rate of biomass loss per hour |
| allowCellOverlap | True | Whether multiple colonies could share a box |
| minSpaceBiomass | 1E-14 | Minimum grams of biomass allowed in a box |
| maxSpaceBiomass | 5E-3 | Maximum grams of biomass allowed in a box |
| defaultVmax | 1 | mmol / gram / hour, max uptake rate or growth rate |
| defaultKm | 0.005 | mM, concentration of half-maximum uptake rate |
| exchangeStyle | Monod Style | Uptake is governed by monod kinetics |
| defaultDiffConst | 5e-6 | cm<sup>2</sup> / s, diffusion constant of resources / metabolites |
| flowDiffRate | 3e-9 | cm<sup>2</sup> / s, diffusion constant of bacterial biomass |
| numDiffPerStep | 10 | How many iterations to solve the diffusion equations<br>per timestep |
| timeStep | 0.1 | Hours, amount of time per timestep |
| Resource concentration | 1e-6 | Mmols of resource initiated in a box |

#### Statistics

The Voronoi areas and other distance metrics (nearest neighbor, summed inverse distances, summed squared inverse distance) were all calculated in R. To find the Voronoi areas of colonies, we used the spatstat package in R and ran the dirichletArea function. After calculation of any distance metric, colonies <5mm from a Petri dish edge were excluded, i.e., the colonies were not included in the analysis, but they did contribute to the distance metric for included colonies.

We used relative metrics to compare yields between treatments and between experiments and simulations. We compared variation in colony yield on different carbon sources by first calculating the coefficient of variation in colony area for each Petri dish, then used a single factor ANOVA to compare the coefficient of variation between experimental treatments. Similarly, to compare experiment to simulation the yields at the end of growth, in units of area for the experiments and grams for the simulations, were divided by the total yield in each Petri dish or simulation area to obtain dimensionless values that could be directly compared. The experimental relative yields were compared with the simulation predictions using mixed-effects linear regression, with the Petri dish as a random factor and the treatment as a fixed factor.

To compare the strength of spatial competition among groups (media type, resource quality), we did an ANOVA followed by Tukey’s Honest Significant Difference multiple comparisons test. The level of replication was the Petri dish. We conducted a t-test to test whether adding acetate to *S. enterica* Petri dishes with glucose altered the strength of spatial competition.

## Results

### The amount of variation in size between colonies on a surface depends on the resources and species identity

We tested whether species and resource identity influenced the variance in colony size within monoculture plates. Approximately 60 cells of either *S. enterica* or *E. coli* were grown on 1% agar Petri dishes with different carbon sources, with 4-8 dishes per treatment. Colony areas were measured using flatbed scanners and custom software (Fig. 1A, see methods for more detail). Within every plate/replicate and treatment we found a range of colony sizes, as seen in the example density plots of the final colony areas in Fig. 1B, suggesting location was important for size variation. Because the *average* colony size differed substantially across treatments (see, for example, Fig. 1B), we used the coefficient of variation of final colony area (standard deviation over mean) within a plate to compare variation in colony size between treatments. Differences in media and species caused large differences in the coefficient of variation across treatment (ANOVA, F(7,32) = 18.9, p = 1.07e-9, Fig. 1C), suggesting that spatial effects were highly context dependent.

### The effects of resource and species identity on the variance in colony size can be predicted with models that pair metabolism and diffusion

We tested whether the observed variations in colony size could be predicted from the interplay of intracellular metabolic mechanisms, diffusion, and colony location, by running simulations that combine genome-scale metabolic modeling with diffusion calculations. Our computational platform, COMETS, uses dynamic flux-balance analysis to predict the growth and metabolic activity of bacteria by identifying the metabolic strategy that maximizes biomass production at each time step (Mahadevan et al. 2002; Harcombe et al. 2014). Biomass and metabolites diffuse to simulate growing colonies and the resource gradients that arise as a result of microbial metabolism. Note that this does not mean that colonies spread by diffusion alone from the center of the colony. Rather, the combined action of diffusion and growth leads to biomass spread (Murray 2002).

Simulations were initiated with resources and colony locations that matched each experimental plate (Fig. 2A). We plotted the relative yields (yield of a colony / total yield on a Petri dish) for simulations against those for experiments (Fig. 2B). We used relative yields because the measurements of interest were the relative differences between colonies on a plate, which can be compared with relative numbers even if the specific yield measurement (area vs biomass) differs. The relative colony sizes in simulations were well correlated with the relative colony sizes in experiments, although the predictive ability of the simulations depended on the treatment (mixed-effects linear regression with subsequent F tests, main effect of simulated yields: F(1,1479) = 2027, p < 2.2e-16, main effect of treatment: F(5,24) = 11, p = 1.3e-5, interaction: F(5,1470) = 85, p < 2.2e-16). Deviations from simulated predictions had a slope < 1, meaning there was more variability in colony size in simulations than in experiments, which occurred and was most pronounced when the carbon resource was a sugar (i.e. glucose or lactose). Below we further explore the deviations caused by growth on sugars.

### Relative colony size is driven by the location of adjacent competitors

To more extensively investigate how location determines colony size we simplified our model to allow faster simulation and abstract species/environment-specific intracellular metabolism.

We replaced the genome-scale metabolic model with a set of differential equations, which describe a reaction-diffusion system in which bacteria grow under Monod kinetics: The growth rate f(B,R) depends on biomass concentration B and resource concentration R. Bacteria and resource spread via diffusion, with the diffusion coefficient for the bacteria *D_B_* being much smaller than that for the resource, *D_R_* (equations 1):

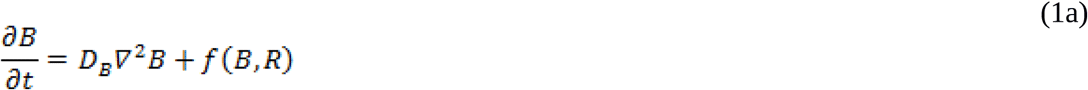

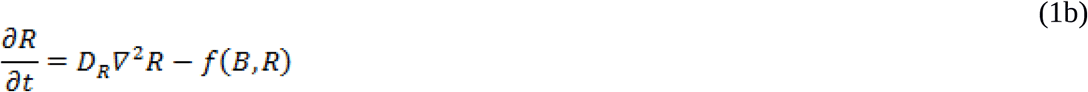

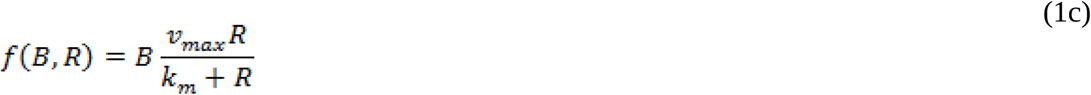

The first term in each of the differential equations describes diffusion (*∇*^2^ is the two-dimensional Laplacian operator.), the second term (the “reaction” term) describes conversion of the resource into biomass which occurs with the same efficiency independent of the growth rate. The maximum growth rate, *v_max_*, is approached as the resource concentration increases, with half-maximum growth rate attained when the resource concentration is equal to the saturation constant *k_m_.*

With this model, we sought a distance-based metric that could predict colony size independently of intracellular details, focusing on metrics that had previous use in the forestry or microbiology literature(Tome & Burkhart 1989; Guillier et al. 2006; Lloyd & Allen 2015). We first simulated conditions that caused strong spatial effects (large variance in colony size), which used a high growth rate (*v_max_* = 1). We tested whether colony size could be best predicted by the distance to the nearest neighbor (Fig. 3A), the sum of the inverse distances to all neighbors (Fig. 3B), the log of the sum of the squared inverse distances (Fig. 3C), or a colony’s Voronoi area, which is the area on the Petri dish which is closer to the focal colony than to any other colony (Fig. 3D)(Okabe et al. 2000). While all metrics were somewhat predictive of colony size, the Voronoi areas had almost perfect prediction.

**Figure 3:**
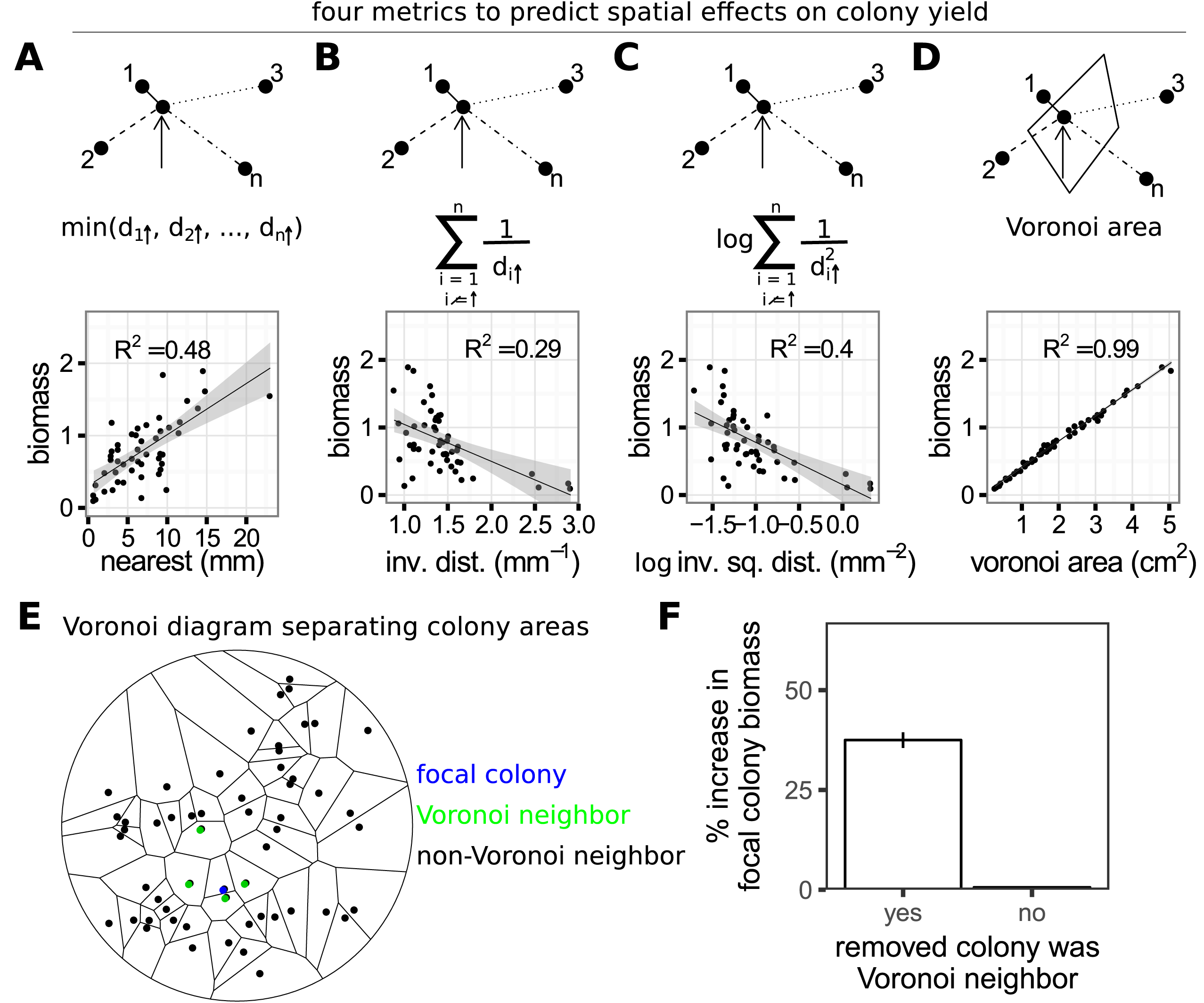
Voronoi diagrams capture the effect of location on yield better than other distance metrics. Four metrics were tested to determine which colonies interact to generate variation in colony size and to what extent. A-D show a cartoon of the measurement and the metric plotted against simulated colony yield (biomass). A) The distance to the closest colony, such that the yield of the focal colony (indicated by the arrow) would be predicted from the distance to colony 1, which is closest, but no other colony would be considered. B) The inverse linear distance to every colony, such that the yield of the focal colony would be predicted by the distance to every colony, with each colony’s influence inversely proportional to its distance. C) Like B, but colonies become quadratically less important as distance increases. D) The territory closest to a colony, described by a Voronoi diagram. Here, the focal colony’s Voronoi area is shown (solid line polygon). A Voronoi diagram divides a plane into areas around colony initiation sites such that all the space in a territory is closer to its enclosed colony than to any other colony, which is accomplished by drawing perpendicular lines half-way through lines connecting a focal colony to Voronoi neighbors. E) A Voronoi diagram drawn for all colony initiation sites on a Petri dish. For a focal colony (blue), its Voronoi neighbors are the green colonies. F) The percent increase in a focal colony’s yield, after removal of a Voronoi or non-Voronoi neighbor. Error bars are standard error.

To further test whether Voronoi neighbors are the primary competitors, we ran “dropout” simulations, in which we repeatedly simulated the same environment (i.e. founding cell locations) but removed a different seed of a colony from each simulation. We then determined the impact that each removal had on focal colony sizes, to test the effect of removing a Voronoi neighbor vs. a non-Voronoi neighbor. Removing Voronoi neighbor colonies had much larger effects than removing non-Voronoi neighbors, Fig. 3F (t.test, p = 5.6e-7). Taken together, this means that colonies that determined territory boundaries (the Voronoi neighbors) played the most important role in causing spatial effects.

Finally, we tested the relative predictive ability of Voronoi areas as spatial effects decreased.

We repeated the metric comparison at increasing resource diffusion coefficients *D*_R_, spanning a range that at the low end is similar to the diffusion coefficient of a large protein such as bovine serum albumin in 1% agar (~1E-7 cm^2^ / s), passing the typical diffusion coefficient for sugars (~5E-6 cm^2^ / s), and reaching past the diffusion coefficient of sodium choride (~1.5E-5 cm^2^ / s) (Schantz & Lauffer 1962). Even as the resource diffusion coefficient increased, causing competition to become more global (described below), the Voronoi areas were still the best predictors of yields (Supp. Fig 1). Therefore, we concluded that the territory closest to a colony, obtained using a Voronoi diagram and determined by the competitors which are in adjacent Voronoi areas, is generally the best predictor for how competition for diffusing resources affects colony size.

### The influence of Voronoi neighbors is determined by the competing effects of resource uptake and diffusion

Our results above show that Voronoi area is the best metric for capturing the effect of location on colony size, so we next used this metric to investigate how diffusion and resource usage parameters influence the strength of spatial effects. To analyze the impact of diffusion and uptake we focused on the slope of the line that is generated when plotting 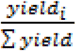 against 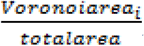, for individual colonies. If each colony has exclusive access to the resources in its territory, the plot should generate a line with a slope of 1. A smaller slope represents increased movement of resources between Voronoi areas. For example, if a plot of 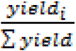 over 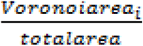 yields a line with a slope of 0.5, a two-fold increase in a colony’s Voronoi area only allows that colony to consume one-fold more resources (since all resources all turned into biomass, independent of rate). In a situation where the slope equals 0, the yield of colonies is largely the same and independent of Voronoi areas (Fig. 4A). Intuitively, a decrease in the slope should occur as competition becomes more global. These interpretations are consistent with the observation that a decrease in the slope, which we deemed the “relative effect of Voronoi area”, goes along with a decrease in the yield coefficient of variation and the R^2 of the plot (Fig. 4B) when varying any parameter (*v_max_*,*k_m_*, R, *D_B_,* or *D_R_*). The correlation between these metrics indicates that parameters that reduce the slope also reduce the amount of variation in colony size and the predictive ability of Voronoi areas.

**Figure 4:**
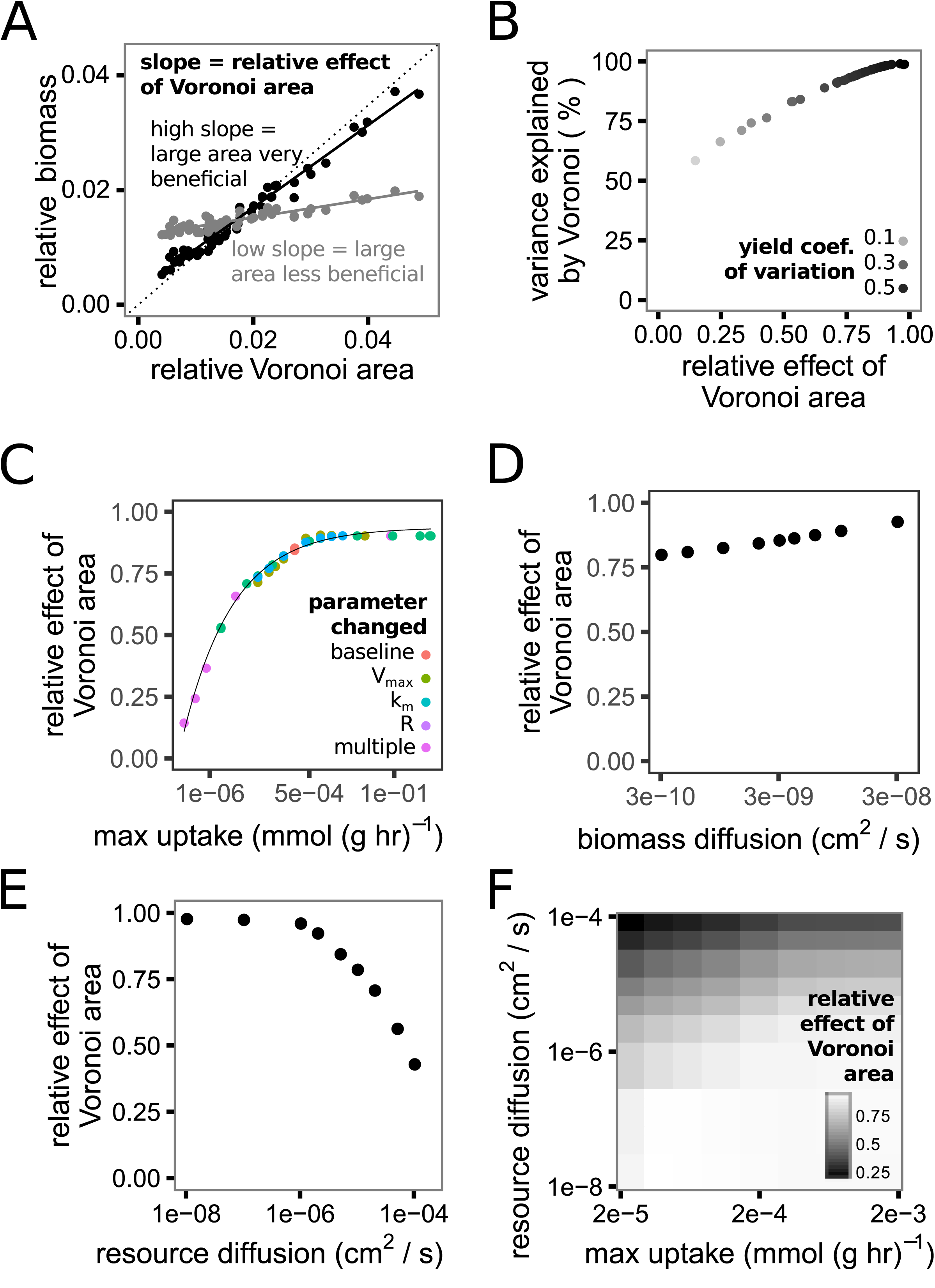
Colony resource uptake rate and resource diffusion determine the relative effect of Voronoi area. A) We quantified the relative effect of Voronoi area by measuring the slope of a line through the standardized colony biomass yields over the colonies’ standardized Voronoi areas. A slope = 1 means that there are strong spatial effects and Voronoi neighbors exert total influence over a focal colony’s yield, whereas a slope = 0 means that all colonies exert similar effects regardless of spatial proximity and therefore competition is global. B) As relative effect of Voronoi area increases, so does the variance explained by the Voronoi areas (100 * R^2^) and the coefficient of variation of colony yield when varying any model parameter. C) The relative effect of Voronoi area increases with the maximum potential per-mass uptake. Changes from the default (baseline) values (Table 1) in maximum growth rate, km, starting resource concentration, or any combination of these parameters (multiple) all have similar effects. D) The relative effect of Voronoi area increases as the biomass diffusion coefficient increases. E) The relative effect of Voronoi area decreases as resource diffusion increases. F) The relative effect of Voronoi area is determined by the balance between the maximum uptake rate of a colony (x axis) and the rate of resource diffusion (y axis).

We next tested what parameters in the model altered the relative effect of Voronoi area. Increasing the rate at which colonies consume nutrients increased the relative effect of Voronoi area. The effects of changing the upper limit on uptake rate (Vmax), the saturation constant (km), or the initial resource concentration (R) all collapsed into a single effect, the maximum uptake rate, by calculating the Monod maximum uptake using Vmax * R / (km + R) (Figure 4C). Increasing the maximum uptake rate caused a fast increase in the relative effect of Voronoi areas until saturating.

We next investigated the influence of biomass and resource diffusion. Increasing the biomass diffusion coefficient (*D_B_*) increased the relative effect of Voronoi area (Fig. 4D). We hypothesize this effect was due to a concomitant increase in the rate at which colonies took up resources that occurred because colonies spread more quickly and were able to reach high resource concentration zones faster. In otherwise identical simulations, faster biomass diffusion did increase the speed at which resources were consumed (Supp. Fig 2).

In contrast to the effect of resource uptake and biomass diffusion, increasing resource diffusion (*D_R_*) *reduced* the relative effect of Voronoi areas. As the resource diffusion coefficient increases, the relative impact of Voronoi neighbors decays (Fig. 4E) likely because diffusion outpaces resource uptake. As a result the impact of Voronoi neighbors is determined by the interplay of nutrient uptake and diffusion (Fig 4 F).

### In laboratory experiments, growth rate predicts the relative effect of Voronoi area

We tested the predicted explanatory power of growth parameters on our laboratory data from Fig. 1. In every treatment, the colony sizes scaled with the colonies’ Voronoi area, as seen in the scatterplots with data from four Petri dishes per treatment shown in Fig. 5A. The relative effect of Voronoi area depended on the species and the environment, Fig. 5B (two-way ANOVA, effect of media: F(4,36) = 40.6, p < 1e-10), no main effect of species, interaction between media:species: F(2,36) = 5.88, p = 0.0062).

**Figure 5:**
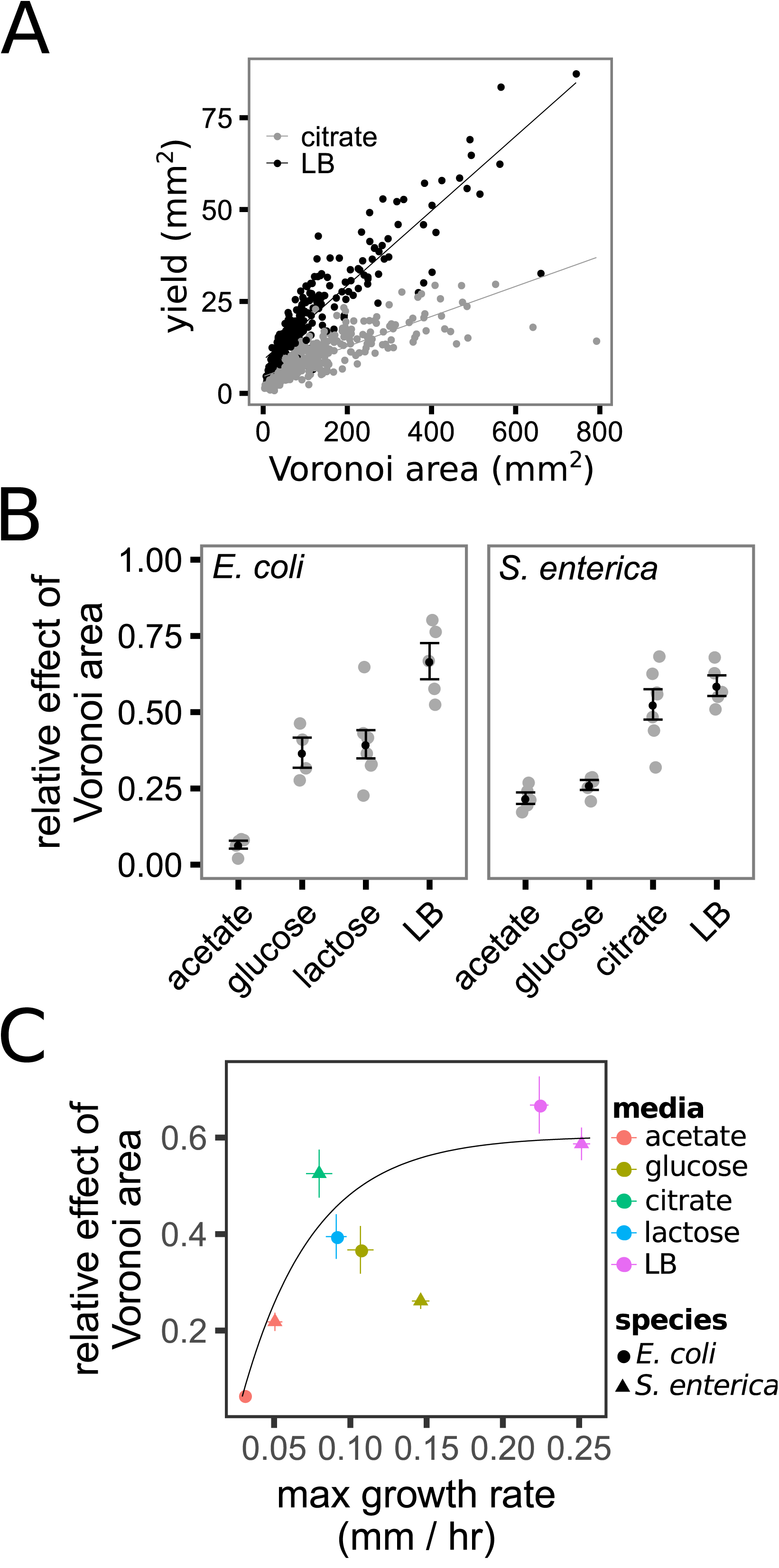
The relative effect of Voronoi area changes between experimental treatments but generally scales with the maximum colony growth rate. A) Scatterplots of the yield of *S. enterica* colonies over their Voronoi areas when grown on citrate or LB media. B) The relative effect of Voronoi area varied from extremely low *(E. coli* on acetate) to high (either species on LB) in experiments. The gray dots are measurements from individual Petri dishes, and the black dots and bars are means and standard error of the means, respectively. C) The relative effect of Voronoi area plotted over the maximum growth rate. The black line is a fit to data, excluding data from *S. enterica* grown on glucose.

In simulations, the maximum growth rate and resource uptake rate were directly proportional, and we showed than increasing the maximum uptake rate increased the relative effect of Voronoi area. In the laboratory data, we measured the maximum growth rate as the increase in diameter over the first three hours after colonies were identified(Palumbo et al. 1971). Media and species each caused significant differences in the maximum growth rate, Fig. 5C (ANOVA, effect of media: F(4,38) = 260, p < 2e-16, effect of species: F(1,38) = 28.2, p = 5.1e-6). Furthermore, in agreement with simulations, the relative effect of Voronoi area in laboratory experiments increased as maximum growth rate increased following a saturating function (Fig. 5C). However, there was the appearance of an outlier: *S. enterica* grown on glucose.

*S. enterica* grown on glucose appeared not to follow the growth rate—Voronoi effect trend (Fig. 5C), and also was the treatment most poorly predicted by the genome-scale metabolic modeling (Fig. 2B). This led us to hypothesize that another biological phenomenon besides competition for diffusing resources was occurring in this treatment. Interestingly, Voronoi areas and metabolic models did a good job of predicting the size of small colonies however, large colonies were routinely smaller than predicted (Fig. 2B). This suggests that large colonies stopped growing before they ran out of resources. *S. enterica* can generate potentially-toxic acetate during growth on glucose, so we hypothesized that acetate accumulation arrested growth of large colonies (Cappuyns et al. 2009; Wolfe 2005; Rhee et al. 2003). To test this hypothesis we grew *S. enterica* on Petri dishes with glucose medium with or without supplemented acetate (each at the same concentration as used in single carbon cultures), reasoning that if acetate was accumulating and causing toxicity, thereby reducing the spatial effects, supplementing acetate would further reduce the spatial effect. Consistent with our hypothesis supplemental acetate (but not growth on acetate alone) reduced the growth of the colonies (Fig. 6A) and the relative effect of Voronoi area (Fig. 6B). Furthermore, genome-scale metabolic modeling (which does not model toxicity) better recapitulated the laboratory data at an earlier time point, when less acetate would have accumulated. This was true for *S. enterica* on glucose, as well as for *E. coli* on glucose or lactose, each of which can cause acetate accumulation (Fig. 6C) (Wolfe 2005).

**Figure 6:**
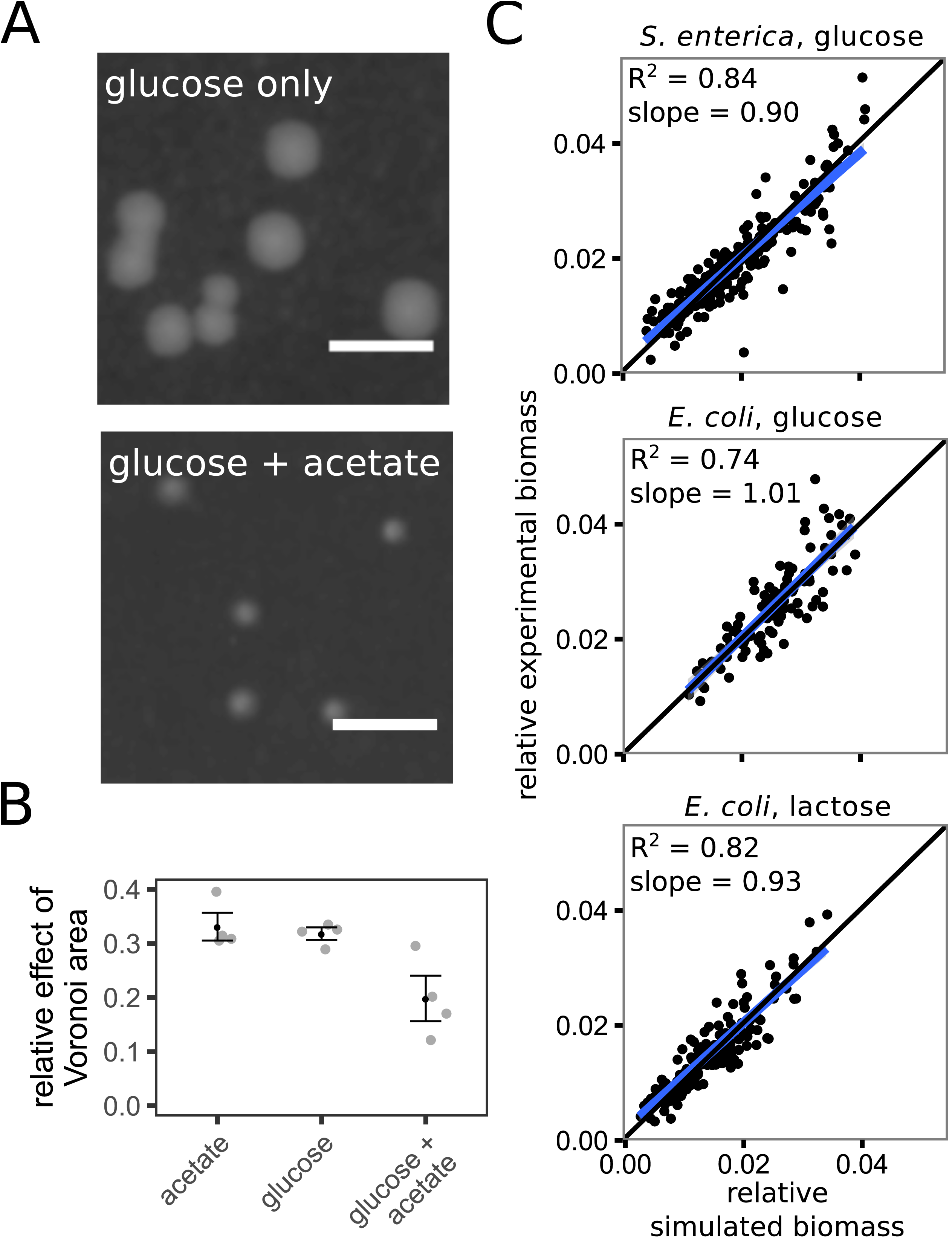
Acetate addition or accumulation reduces the relative effect of Voronoi area. A) *S. enterica* colonies grown on Petri dishes with glucose alone or with acetate added. The scalebar = 2.5mm. B) The relative effect of Voronoi area with only acetate, only glucose or a combination of the two. The gray dots are measurements from individual Petri dishes, and the black dots and bars are means and standard error of the means, respectively. C) The simulations from Fig. 2B, but with the simulated results taken from 40 hours into the simulation rather than at the end of the experiment (150 hrs). Acetate accumulation is not predicted to occur until 40 hours.

## Discussion

Understanding the quantitative way that spatial proximity affects interactions between bacterial colonies will allow us to better understand and manage microbial ecosystems. We showed that while species identity and the resources present caused variations in the effect of colony proximity, we could nevertheless predict much of this variance using models accounting for metabolism and diffusion. Spatially-dependent variation in colony size was largest on media which promoted a fast growth rate. Specific colony sizes were best predicted using Voronoi diagrams, which means that the differential colony growth was primarily caused by adjacent competitors (the “Voronoi neighbors”) (Okabe et al. 2000). The specific influence of these Voronoi neighbors was determined by a balance between the rate at which colonies took up resources, and resource diffusion. High uptake rates increased the importance of Voronoi neighbors and caused greater spatially-dependent variance in colony size. This general ecological relationship held across species and environments and therefore serves as a useful null model from which to predict spatial effects caused by resource competition. We demonstrated the utility of this null model: experiments in which Voronoi neighbors had less influence than would be predicted from growth rate led us to suggest that the toxicity of organic acid byproducts plays an important role in limiting colony size when bacteria are growing on sugars. In summary, we provide an experimentally and theoretically grounded understanding of how location interacts with metabolism and diffusion to influence microbial interactions.

We found that the impact of location on microbial growth was strongly influenced by both species and resource identity. This finding highlights the fact that the impact of spatial structure is context-dependent. While a dichotomy between structured and unstructured environments has value (Kim et al. 2008; Kerr et al. 2002; Chao & Levin 1981), it is important to realize that the effect of structure can change dramatically in different environments (Allen et al. 2013). As we strive to understand interaction strengths in natural microbial communities and design spatially-structured ecosystems for technological applications, it will be vital to incorporate context-dependent effects of location.

Encouragingly, spatially-explicit, genome-scale metabolic models were able to predict much of the variation in colony size by modeling the interaction between diffusion and intracellular metabolism. This suggests that with models created from sequence data we will be able to quantitatively predict metabolic microbial interactions in complex, spatially structured environments. High-throughput methods to generate models from sequence data are improving(Feist et al. 2009), and therefore spatially explicit tools such as COMETS will be increasingly useful to generate quantitative predictions of the effect of location on growth and microbial interactions. As we discuss below, the accuracy of the predictions will be strongly influenced by realized uptake rates. The upper-limit of uptake (Vmax) must be defined in the model but will often be difficult to infer from sequence data. However, in our simulations we left the upper-limit for all metabolites set at a canonical value of 10 mmol / gram dry weight / hr (Harcombe et al. 2014) and still achieved a good match between simulation and experiment. This suggests that colony-level uptake rates depended more on the stoichiometry of biomass production on each carbon source than on Vmax. Therefore, accurate predictions of stoichiometry, obtainable from sequence data, may lead to good approximations of the rate at which growing colonies consume resources even with inexact limits on uptake rates.

Voronoi diagrams did substantially better at explaining colony variance than did any of the other metrics that we tried. Dropout simulations show that colonies outside the Voronoi neighbors have very little influence on variation in colony size. Interestingly, this suggests that for competition the arrangement of nearby colonies matters more than the specific proximity of those colonies. One caveat of our findings is that all of our colonies started to grow at roughly the same time. Had bacteria colonized the plate at different time points, or varied dramatically in lag time, Voronoi diagrams would likely explain less of the variance. Indeed differences in lag time have been shown to influence competition for physical space (Lloyd & Allen 2015), however, lag time had exceedingly little impact on colony variance in our experiments (data not shown). Additionally, we only looked at interactions with conspecific colonies. We expect that predicting interactions between species with different resource uptake rates will require weighted Voronoi diagrams. Finally, in future work it will also be interesting to investigate how Voronoi diagrams fair in three-dimensional ecosystems, such as lung infection models (Connell et al. 2014).

A balance of resource uptake and diffusion determined the extent to which competitive interactions between bacterial colonies were localized. Any parameter that increased uptake increased the influence of Voronoi neighbors (up to a limit), but increasing resource diffusion mitigated this effect. This means that both uptake and diffusion must be considered to determine the extent to which interactions are local. One might expect non-localized interactions in aquatic environments due to the low viscosity of the media, but if liquid is static, and resource-rich, the bacteria nevertheless may interact primarily with Voronoi neighbors. Conversely, in structured environments where one might expect highly localized interactions, if nutrient uptake is slow enough, resource diffusion may overpower resource uptake and globalize the interactions. It is important to note that decreasing the extent to which competition is local is not equivalent to decreasing competition. The average colony size and total biomass on a plate are equivalent whether competition is local or global (assuming all resources are consumed). However, if the balance of uptake and diffusion cause interactions to be local, spatial location matters, and some colonies will grow much larger than others.

The balance between resource uptake and diffusion provides a null model for colony variance given resource competition in a spatial environment. If colony growth is mediated by competition for relatively slowly diffusing resources, then colony size should correlate with Voronoi area. Departures from this null expectation can help identify circumstances in which other biological interactions are occurring. In our experiments, colony size of *S. enterica* correlated poorly with Voronoi area when glucose was the carbon source, despite a fast rate of growth. Here, waste accumulation appears to have stunted the growth of large colonies. Both *E. coli* and *S. enterica* produce organic acids as a byproduct of growth on sugars, and acetate toxicity can hinder growth(Wolfe 2005; Vandenbergh 1993). This toxicity reduced the relative importance of location. As production of toxic byproducts is common in the microbial world, it will be interesting to further investigate how toxicity influences spatial patterns. More broadly, the detection of toxicity in our system serves as an example of how quantitative analysis can aid in the identification of species interactions. Different biological phenomena likely cause specific departures from the null expectations. For instance, we hypothesize that mutualistic interactions between colonies will cause spatial effects, but of opposite direction to competition, in which colonies in smaller Voronoi areas have better success. Further research will be aimed at finding signatures of these and other biological phenomena.

A quantitative understanding of how location mediates microbial interactions has important consequences for understanding and harnessing microbial evolutionary ecology. It is well established that spatial structure matters, can alter the interactions between microbes(Nadell et al. 2016) and plays a critical role in determining health outcomes(Stacy et al. 2016). Quantifying how space mediates interactions will allow for more rigorous understanding of community composition, and improve prediction of dynamics such as competitive exclusion. Further, understanding organisms’ interaction strengths is critical for understanding the evolution of microbial traits. For example, it was recently demonstrated that the level of antibiotic secretion can be explained by the relative strength of interaction with sensitive and resistant competitors (Gerardin et al., 2016). As technology which allows for fine-scale placement of cells matures (Xu et al. 2004; Ferris et al. 2013; Connell et al. 2013), we can create spatial arrangements that maximize selection of competitive phenotypes of interest. As we strive to move beyond descriptions of microbial diversity to explanations and management of diversity it will be critical to develop quantitative understanding of microbial interactions.

## Acknowledgements

The authors thank S. Zhuang for advice on the scanning technique and the UMN theory group for useful discussions. J. Chacón was funded by the Biocatalysis Initiaitive through UMN.

---

**Supplementary Figure 1: Voronoi areas are the best predictors of yield even as the system becomes more global** The amount of yield variance explained by each metric as the resource diffusion rate is increased. The colony layout is the same as for the data in Fig. 3. Regardless of the diffusion rate, Voronoi areas are the superior metric.

**Supplementary Figure 2: Total consumption of the ecosystem’s resources occurs more quickly as the biomass diffusion coefficient increases** The time step at which 99.9% of the simulated ecosystems’ resources were consumed, plotted over the biomass diffusion constant. All other parameters were held constant. The data is from the same simulations as shown in Fig. 4D.

